# Identifying transposon insertions in bacterial genomes through nanopore sequencing

**DOI:** 10.1101/765545

**Authors:** David A. Baltrus, John Medlen, Meara Clark

**Affiliations:** School of Plant Sciences, University of Arizona, Tucson AZ, USA; School of Animal and Comparative Biomedical Sciences, University of Arizona, Tucson AZ, USA

## Abstract

Transposon mutagenesis is a widely used tool for carrying out forward genetic screens across systems, but in some cases it can be difficult to identify transposon insertion points after successful phenotypic screens. As an alternative to traditional methods, we report on the efficacy of using an Oxford Nanopore’s MinION to identify transposon insertions through whole genome sequencing. We also report experiments using CRISPR-Cas to selectively target regions of the genome where a transposon has integrated. Our experiments provide a framework for understanding the efficiency of such techniques for carrying out forward genetic screens and point towards the ability to use CRISPR-based sequence capture to identify the insertion of particular regions of DNA across all genomes, which may enable Tn-Seq experiments using Nanopore based sequencing.

## Introduction

Forward genetic experiments (whereby phenotypes of interest are first screened and then the causative mutations of interesting phenotypes are identified) have broadly contributed to building the foundation of modern microbial genetics (Shuman & Silhavy, 2003). Among these, experiments that utilize transposon insertions have been particularly useful in generating the causative mutations for screens as well as providing a means to identify which region of the genome was disrupted (Barquist, Boinett & Cain, 2013; Chao et al., 2016). Since identification of transposon insertion sites can sometimes be challenging, our goal here is to investigate the use of Nanopore based sequencing to interrogate the positions of transposon insertions in bacterial genomes.

The design of transposon mutagenesis experiments can be significantly limited by the study systems of interest (e.g. types of antibiotics available for selection, effectiveness of transposon mutagenesis within each background, etc…), and the further away from established model systems one goes the more challenging such experiments can be. In some cases, significant steps are required to engineer transposon sequences and delivery systems to ensure experimental success. Moreover, even when transposon mutagenesis works well, such experiments can be limited by availability or effectiveness of tools like PCR primers to categorize insertion points across systems. To aid these experiments, tools like modified Tn5 vectors have been developed to streamline categorization of insertion points through chromosomal digestion with restriction enzymes followed by ligation and recapture cloning (Larsen et al., 2002), but successful implementation of these protocols can be time consuming and dependent on precise molecular techniques. Alternatively, a variety of transposons have been recently engineered which can be sequenced and screened in a high throughput way through TnSeq (Barquist, Boinett & Cain, 2013; Chao et al., 2016), but technological, monetary, and computational requirements of these approaches could limit their adoption and use.

Identification of transposon insertions sites through sequencing of whole bacterial genomes using an Oxford Nanopore MinION could empower labs to carry out cost and time efficient forward genetics experiments across many different systems (Jain et al., 2016). Importantly, although Nanopore based sequencing has already been widely implemented, adoption of Nanopore only sequencing strategies remain limited by relatively high per-read error rates compared to other platforms (Tyler et al., 2018). However, identification of transposon (or any DNA with known sequence) insertion points should not be greatly affected by error rates so long as parts of the read are alignable. Therefore, querying the relatively long reads generated by MinION sequencing for transposon insertions should provide a rapid and straightforward means to identify insertion positions. Furthermore, techniques have recently been developed that allow for targeted sequence capture of genomic regions through the use of CRISPR-Cas9 enrichment of genomic DNA (Gilpatrick et al., 2019). These approaches should work well for identifying transposon insertions because the transposon sequences are known and are unchanged regardless of where in the genome they integrate. All cost-benefit calculations for implementing this technology depend upon the ability to sequence and identify transposon insertions from long reads as well as the efficiency of calling reads that have been multiplexed, and to this point there is currently little data vetting the positives and negatives of identifying transposon insertions within bacterial genomes using the MinION.

Here we evaluate the ability of multiplexing genomes across two different sequencing kits in order to identify transposon insertions and compare this to the ability of CRISPR-Cas9 targeted enrichment to identify transposon insertions. Since the efficacy of these approaches ultimately depends upon identifying the transposon signal amongst the noise of all of the other genomic sequences, we highlight critical bottlenecks in the process of using Nanopore sequencing for transposon based forward genetic screens and areas in which subtle tweaks and improvements could make significant differences in the utility and cost efficiency of these approaches. Lastly, we report the use of CRISPR-Cas9 genomic enrichment to identify transposon insertion points within bacterial genomes. The success of these approaches could open the door to effective TnSeq experiments across systems and regardless of transposon identity using Nanopore based sequencing.

## Methods

### Strains for Sequencing

We used derivatives of five different strain backgrounds to carry out these sequencing experiments: *Ensifer adhaerens* Casida A (DBL796), *Luteibacter* sp. 9143 (DBL564), *Pseudomonas* spp. Leaf58 (DBL1616), *Pseudomonas putida* KT2440 (DBL1620), and *Pseudomonas stutzeri* 28a24 (DBL494). Details underlying genotypes, transposon mutagenesis, and phenotypic screening of specific strains sequenced as a part of experiments presented here can be found at https://doi.org/10.6084/m9.figshare.8795144.v2.

### Isolation of Genomic DNA

Each strain was streaked to single colonies on either Lysogeny Broth (LB, *Luteibacter* and *Ensifer*) or King’s Medium B (KB, *Pseudomonas*) agar plates supplemented with antibiotics specific to each transposon (100ng/uL gentamicin for pMar2xT7 derivatives in *E. adhaerens* and 25ng/uL for pRL27 derivatives). A single colony of each strain was then picked to a test tube containing 2mL liquid media (again, LB or KB) supplemented with appropriate antibiotics and grown overnight at 27°C in a shaking incubator at 220rpm. The next day, genomic DNA of each culture was extracted using a Promega Wizard Kit as per manufacturer’s protocols and including RNAse addition. Genomic DNA was prepared in the same way but independently for strains across all three sequencing experiments (RAPD, LSK109, CRISPR-Cas9 enriched LSK109). Genomic DNA was quantified by Qubit with Broad sensitivity range kit, and roughly 100ng of each sample of genomic DNA was run and visualized in a 1% agarose gel to gauge overall size and fragmentation of genomic DNA (gel pictures available at https://doi.org/10.6084/m9.figshare.8795144.v2).

### Rapid Kit Based Sequencing

400ng of DNA from each of the 12 strains was added to individual reactions using each barcode from the RAPD barcoding kit (SQK-RPB004) per the manufacturer’s instructions. *Ensifer adhaerens* strains were given barcodes 1-5, *Pseudomonas* sp. Leaf58 strains were given barcodes 6-8, *Luteibacter* sp. 9143 strains were given barcodes 9 and 10, with *P. stutzeri* barcode 11 and *P. putida* barcode 12. After cleanup with AmpPure beads, all reactions were pooled in equal volumes and prepped for sequencing on a MinION. Raw reads arising from the “fast5_pass” folder were concatenated together into a file called RAPD_tn_final.fast5.tgz, which is publicly available at the following link: https://de.cyverse.org/dl/496A93AE-ACAB-4112-8EBD-8CD01D3CFF82

Basecalling took place in real time using Guppy version 2.0.10, and only reads passing initial quality checks (QC) during the run (and which were therefore placed into the “fastq_pass” folder) were used for subsequent steps. Reads were demultiplexed using qcat version 1.0.7, and these demultiplexed reads were used for all downstream analyses.

### Ligation Kit Based Sequencing

A separate genomic extraction was carried out, as above and starting with a single colony, for each of the 12 focal strains. 1ug of DNA from the strains was barcoded for sequencing on a MinION using the NBD-003 kit and per the manufacturer’s instructions. Strains were barcoded in the same order as with the RAPD Kit (1-12), with the exception that barcodes 8 and 9 were switched compared to the RAPD kit preparation. After cleanup with AmpPure beads, all reactions were pooled in equal volumes and prepped for sequencing on a MinION using the LSK-109 kit. Raw reads arising from the “fast5_pass” folder were concatenated together into a file called LSK109_tn_final.fast5.tgz, which is publicly available at the following link: https://de.cyverse.org/dl/58DAD7C1-89EC-475F-ACE9-7E90095500B9

Basecalling took place in real time using Guppy version 2.0.10, and only reads passing initial quality checks (QC) during the run (and which were therefore placed into the “fastq_pass” folder) were used for subsequent steps. Reads were trimmed and demultiplexed using qcat version 1.0.7, and these demultiplexed reads were used for all downstream analyses.

### Targeted Genome Sequencing Using CRISPR-Cas

2.5ug of genomic DNA from a separate genomic DNA extraction of *P. stutzeri* (DBL494) was dephosphorylated following instructions from a protocol distributed by Oxford Nanopore and referenced in Timp et al. After this point, genomic DNA was mixed in a reaction using CRISPR-Cas9 and 4 different crRNA oligos designed to target the Mar2xT7 transposon (sequence of the transposon is available at https://doi.org/10.6084/m9.figshare.8795144.v2).

The sequences of these four oligos were:
pMar2xT7-3) 5’ AlTR1/UAACAUCAAACAUCGACCCAGUUUUAGAGCUAUGCU/AlTR2 3’
pMar2xT7-4) 5’ AlTR1/CUUGAGGAGAUUGAUGAGCGGUUUUAGAGCUAUGCU/AlTR2 3’
pMar2xT7-6) 5’ AlTR1/AGAACCUUGACCGAACGCAGGUUUUAGAGCUAUGCU/AlTR2 3’
pMar2xT7-7) 5’ AlTR1/UUACGGUGACGAUCCCGCAGGUUUUAGAGCUAUGCU/AlTR2 3’

After crRNA targeting, genomic fragments were prepped and sequenced an Oxford Nanopore MinION using kit LSK109. Basecalling took place in real time using Guppy version 2.0.10, and only reads passing initial quality checks (QC) during the run (and which were therefore placed into the “fastq_pass” folder) were used for subsequent steps. Raw reads arising from the “fast5_pass” folder were concatenated together into a file called CRISPR_tn_final.fast5.tgz, which is publicly available at the following link: https://de.cyverse.org/dl/420E53F6-51EC-4F07-B304-6D81F81AA136

### Identification of Total Number of Reads Containing Transposon Sequences

After demultiplexing, reads arising from both the RAPD kit based protocol and the ligation kit based protocol were characterized using a standard pipeline. First, transposon sequences arising from either pMar2xT7 or the modified Tn5 from pRL27 were used as queries against reads from each of the strains of interest (with query sequence matching the transposon within that particular genome) using Blastn (Camacho et al., 2009) with largely default parameters (except evalue 1e-5, outfmt 6). Sequences used to query each are found in a .fasta file available at https://doi.org/10.6084/m9.figshare.8795144.v2. Only hits for the transposon of interest were used in further analyses. A custom perl script, available at https://doi.org/10.6084/m9.figshare.8795144.v2, called ‘FastaGrabber.pl’ was used to select reads containing possible transposon sequences from sequencing runs for each of the strains of interest. These reads were then used as a query sequence against the relevant genome sequence for each isolate using Blastn and largely with default parameters (except evalue 1e-5, outfmt 6, max_hsps 1) with two exceptions. Genome sequences are found on Genbank with accessions: *Ensifer adhaerens* (GCA_000697965.2), *Luteibacter* sp. 9143 (GCA_000745235.1), *Pseudomonas stutzeri* 28a24 (GCA_000590475.1), and *Pseudomonas* sp. Leaf58 (GCA_003627215.1). In the cases of DBL1620 and DBL494, the sequence of pMPPla107 was the target sequence for this Blastn search because it was known that the transposon was contained on plasmid pMPPla107 (Genbank accession of pMPPla107 is NZ_CP031226.1). Positions of these Blastn results were then compared, and all reads mapping within the same 20kb in the target assembly (across both the RAPD and LSK109 libraries) were counted as containing transposon insertions. Transposon mapping results from these comparisons using Blastn were equivalent to those acquired using minimap2 (data not shown).

Reads arising from targeted genome sequencing using CRISPR-Cas against DBL494 were aligned against the sequence of DBL494 (the chromosome of *P. stutzeri* 28a24 and megaplasmid pMPPla107) using minimap2 (Li, 2018) with default parameters.to generate a .sam file. Samtools was then used to generate .bam and sorted .bam files, with the sorted .bam file formatted using BedTools (genomecov -bg) to generate the resulting coverage file. All files are available at https://doi.org/10.6084/m9.figshare.8795144.v2. We then used Circos (Krzywinski et al., 2009) to visualize coverage levels of these reads against the genome of strain DBL494 (containing the chromosome of *P. stutzeri* 28a24 and megaplasmid pMPPla107).

### Identification of Insertion Sites for Transposon Sequences

To identify particular insertion sites for the transposon within each genome, single reads where the transposon sequence of interest was flanked by genomic DNA (as judged by the blastn results) were inspected by hand. Nucleotide sequences flanking these transposon sequences were then aligned to each relevant genome sequence of interest using Geneious Prime 2019.2.1 (www.geneious.com) and default parameters. Insertion sites for each of the strains can be found in a spreadsheet at https://doi.org/10.6084/m9.figshare.8795144.v2.

### Confirmation of Transposon Insertion Sites in Luteibacter Genomes

We used a basic ligation-recapture protocol to identify sites of insertion for the modified Tn5 from pRL27 that transposed into each *Luteibacter* genome (Larsen et al., 2002). Briefly, genomic DNA was isolated from each strain using a Wizard Kit (Promega), and was digested with BamHI. Fragments were then ligated back together into covalent circles, which were then transformed through electroporation into a *pir*+ version of *E. coli* strain S17. Since the modified Tn5 from pRL27 contains both kanamycin resistance and an *ori*R6K origin of replication, genomic fragments that contain this transposon will replicate as plasmids within *pir*+ strains of *E. coli*. Lastly, we isolated plasmids from *E. coli* strains arising from each of these two reactions and sequenced plasmids using primers 13-2 and 17-1 to confirm genomic regions that flanked each transposon. Sequencing (.abi) files from each of these reactions are available at https://doi.org/10.6084/m9.figshare.8795144.v2.

## Results and Discussion

### Identification of transposon insertion sites through whole genome sequencing

Results and metrics from both RAPD and LSK109 sequencing runs can be found in Tables 1 and 2, respectively. Since roughly the same amount of total DNA was sequenced by each kit and the read lengths arising from sequencing using each kit are equivalent, comparison of these tables provides a useful metric for evaluating the ability to identify transposon insertion points from these methods. To further streamline comparisons we have calculated the % of transposon informative reads arising from each multiplexed strain library, which provides an estimate for the number of reads that can be used specifically to identify transposon insertions, and provide these numbers in the tables along with other metrics such as the N50 of the reads lengths for each strain.

**Table 1.** Reads Arising From RAPD Kit Based Sequencing

| Bar Code | Strain | Transposon | Genome Size (bp) | Reads (qcat) | Bases (bp) | Read N50 | Number of Reads Containing Genome and Transposon Sequence | Number of Total Blast Hits To Transposon Query | Percent of Transposon Informative Hits |
| --- | --- | --- | --- | --- | --- | --- | --- | --- | --- |
| 1 | Ensifer adhaerens Casida A | pMar2xT7 | 7,267,502 | 8,456 | 32,002,330 | 6690 | 10 | 14 | 0.12% |
| 2 | Ensifer adhaerens Casida A | pMar2xT7 | 7,267,502 | 10,552 | 43,860,296 | 7293 | 9 | 9 | 0.09% |
| 3 | Ensifer adhaerens Casida A | pMar2xT7 | 7,267,502 | 2,786 | 11,541,859 | 7637 | 1 | 1 | 0.04% |
| 4 | Ensifer adhaerens Casida A | pMar2xT7 | 7,267,502 | 4,291 | 17,611,918 | 7155 | 6 | 20 | 0.14% |
| 5 | Ensifer adhaerens Casida A | pMar2xT7 | 7,267,502 | 4,156 | 18,845,262 | 8094 | 8 | 11 | 0.19% |
| 6 | Pseudomonas sp. Leaf58 | pRL27 | 6,337,121 | 43,471 | 92,827,094 | 3567 | 32 | 36 | 0.08% |
| 7 | Pseudomonas sp. Leaf58 | pRL27 | 6,337,121 | 13,281 | 42,723,957 | 5597 | 12 | 12 | 0.09% |
| 8 | Pseudomonas sp. Leaf58 | pRL27 | 6,337,121 | 4,212 | 12,524,800 | 5425 | 6 | 7 | 0.14% |
| 9 | Luteibacter sp. 9143 | pRL27 | 4,625,266 | 22,879 | 83,299,299 | 6673 | 21 | 26 | 0.09% |
| 10 | Luteibacter sp. 9143 | pRL27 | 4,625,266 | 6,041 | 23,420,139 | 6746 | 7 | 7 | 0.12% |
| 11 | Pseudomonas stutzeri 28a24 + pMPla107_494 | pMar2xT7 | 5,703,249 | 27,589 | 177,256,767 | 11559 | 39 | 40 | 0.14% |
| 12 | Pseudomonas putida KT2440 + pMPla107_H4 | pRL27 | 7,153,762 | 23,445 | 110,315,944 | 8693 | 15 | 516 | 0.06% |
| None Identified |  |  |  | 221,810 | 886,739,766 | 7326 |  |  |  |
| Totals |  |  |  | 392,969 | 1,552,969,431 |  |  |  |  |

**Table 2.** Reads Arising From LSK109 Kit Based Sequencing

| Bar Code | Strain | Transposon | Genome Size (bp) | Reads (qcat) | Bases (bp) | Read N50 | Number of Reads Containing Genome and Transposon Sequence | Number of Total Blast Hits To Transposon Query | Percent of Transposon Informative Hits |
| --- | --- | --- | --- | --- | --- | --- | --- | --- | --- |
| 1 | Ensifer adhaerens Casida A | Mar2xT7 | 7,267,502 | 16,505 | 97,312,007 | 12330 | 13 | 13 | 0.08% |
| 2 | Ensifer adhaerens Casida A | Mar2xT7 | 7,267,502 | 33,730 | 60,417,009 | 2629 | 7 | 9 | 0.02% |
| 3 | Ensifer adhaerens Casida A | Mar2xT7 | 7,267,502 | 14,439 | 80,941,296 | 12669 | 10 | 10 | 0.07% |
| 4 | Ensifer adhaerens Casida A | Mar2xT7 | 7,267,502 | 37,245 | 144,635,155 | 5777 | 35 | 36 | 0.09% |
| 5 | Ensifer adhaerens Casida A | Mar2xT7 | 7,267,502 | 26,521 | 116,831,016 | 6466 | 17 | 18 | 0.06% |
| 6 | Pseudomonas sp. Leaf58 | pRL27 | 6,337,121 | 29,310 | 185,097,035 | 11019 | 38 | 38 | 0.13% |
| 7 | Pseudomonas sp. Leaf58 | pRL27 | 6,337,121 | 15,470 | 98,660,191 | 13281 | 13 | 15 | 0.08% |
| 8 | Luteibacter sp. 9143 | pRL27 | 4,625,266 | 31,078 | 123,352,576 | 5808 | 23 | 23 | 0.07% |
| 9 | Pseudomonas sp. Leaf58 | pRL27 | 6,337,121 | 22,176 | 100,539,904 | 7613 | 18 | 22 | 0.08% |
| 10 | Luteibacter sp. 9143 | pRL27 | 4,625,266 | 8,689 | 48,371,097 | 9305 | 13 | 14 | 0.15% |
| 11 | Pseudomonas stutzeri 28a24 + pMPla107_494 | Mar2xT7 | 5,703,249 | 11,671 | 66,960,491 | 12344 | 14 | 15 | 0.12% |
| 12 | Pseudomonas putida KT2440 + pMPla107_H4 | pRL27 | 7,153,762 | 4,460 | 16,402,247 | 6586 | 2 | 84 | 0.04% |
| None Identified |  |  |  | 16,190 | 75,042,870 |  |  |  |  |
| Totals |  |  |  | 267,484 | 1,139,520,024 |  |  |  |  |

The RAPD kit is a relatively straightforward and fast means to prepare genomic DNA for sequencing on an Oxford Nanopore MinION, and takes advantage of transposon tagmentation of the DNA to incorporate adaptors enabling sequencing on a flowcell. This kit requires the least number of steps to prepare genomic DNA for sequencing, with samples ready to load on a MinION in as little as 15 minutes. The RAPD kit also currently allows for sequence multiplexing of up to 12 libraries at a time on one flowcell. Across the 12 genomes sequenced, we generated a total of 381,221 reads using this method, for a total of roughly 1.5Gb of data using the RAPD. We then used qcat to demultiplex the barcodes and separate out reads into the twelve different libraries for further identification of transposon insertion sites. In our hands in these libraries, a majority of reads (221,810 or 57% of reads and 887Mb total) were not able to be assigned to particular barcodes. Therefore, we inherently see a loss of information because, outside of other means or through logical deduction, we are unable to definitively assign transposon insertions to genomes for this bin. Secondly, we note that even though we started with roughly the same amount of DNA for each library, there is significant variance in terms of how much depth is actually captured and assigned to each library. Overall, the number of transposon containing reads to total reads ranged from 0.03-0.19% in each barcode, and the number of transposon informative reads appears to scale with the total number of reads sequenced (1:2,786 reads in barcode 3 to 39:27,589 in barcode 11). There was no clear relationship between read length and ability to identify transposon insertions, as the number of transposon informative reads from the library with the lowest N50 (32 reads, N50=3567, barcode 6) was equivalent to the number found from the library with the highest N50 (39 reads, N50=11,559, barcode 11). Furthermore, we have confirmed that this method provides an accurate readout of transposon insertion points, as our results match those obtained for two of the Luteibacter strains through traditional ligation-recapture protocols.

Using a ligation based kit, we generated a total of 250,959 reads for a total of roughly 1.1Mb of data, both metrics which are roughly equivalent to our results with the RAPD kit. In contrast to the RAPD kit, we were able to attribute ~93% of reads to one of the 12 strains using qcat and therefore were able to recover a significantly higher number of reads than when using the RAPD kit. As with the RAPD kit, we found that the percentage of transposon informative reads using LSK109 ranged from 0.02 to 0.14% of the total reads for each library. Importantly, we found that the N50 of each sequencing library was roughly equivalent to that of the RAPD kit, which suggests that initial DNA extraction (which was carried out using a Promega Wizard kit in both cases) is the main determinant of read lengths in our hands.

### Identification of transposon insertion sites through whole genome sequencing using CRISPR-Cas Targeted Enrichment

Protocols have recently been developed that enable targeted sequencing of genomic regions of interest through the use of *in vitro* CRISPR-Cas targeting by custom oligos followed by sequencing using a ligation based kit. Successful enrichment that enables identification of transposon insertion sites with much lower levels of sequencing coverage than either of the other methods would facilitate processing of more strains through a single flowcell. Key for this method, however, is understanding the levels of sequencing noise compared to reads that are informative for transposon insertion identification. For this experiment, a total of 5887 reads (totalling 27,488,682 bp) were sequenced. Roughly 1 in 4 of these reads mapped to the region on megaplasmid pMPPla107 identified by other methods as the transposon insertion within this strain. Looking closer, low levels of coverage (between 1-10x depth) were sequenced by this methods throughout both the chromosome and the megaplasmid (Figure 1). However, coverage significantly increased within the 50,000bp surrounding the identified insertion point averaged (to 111x). Moreover, coverage levels continued to improve within the surrounding 1000/500/100/50bp (676x/699x/727x/734x) surrounding the proposed insertion site (Figure 1, coverage file available at https://doi.org/10.6084/m9.figshare.8795144.v2). Therefore, coverage levels around the identified insertion point are roughly 100 fold higher than the background levels of sequencing coverage using this targeted enrichment method.

**Figure 1.**
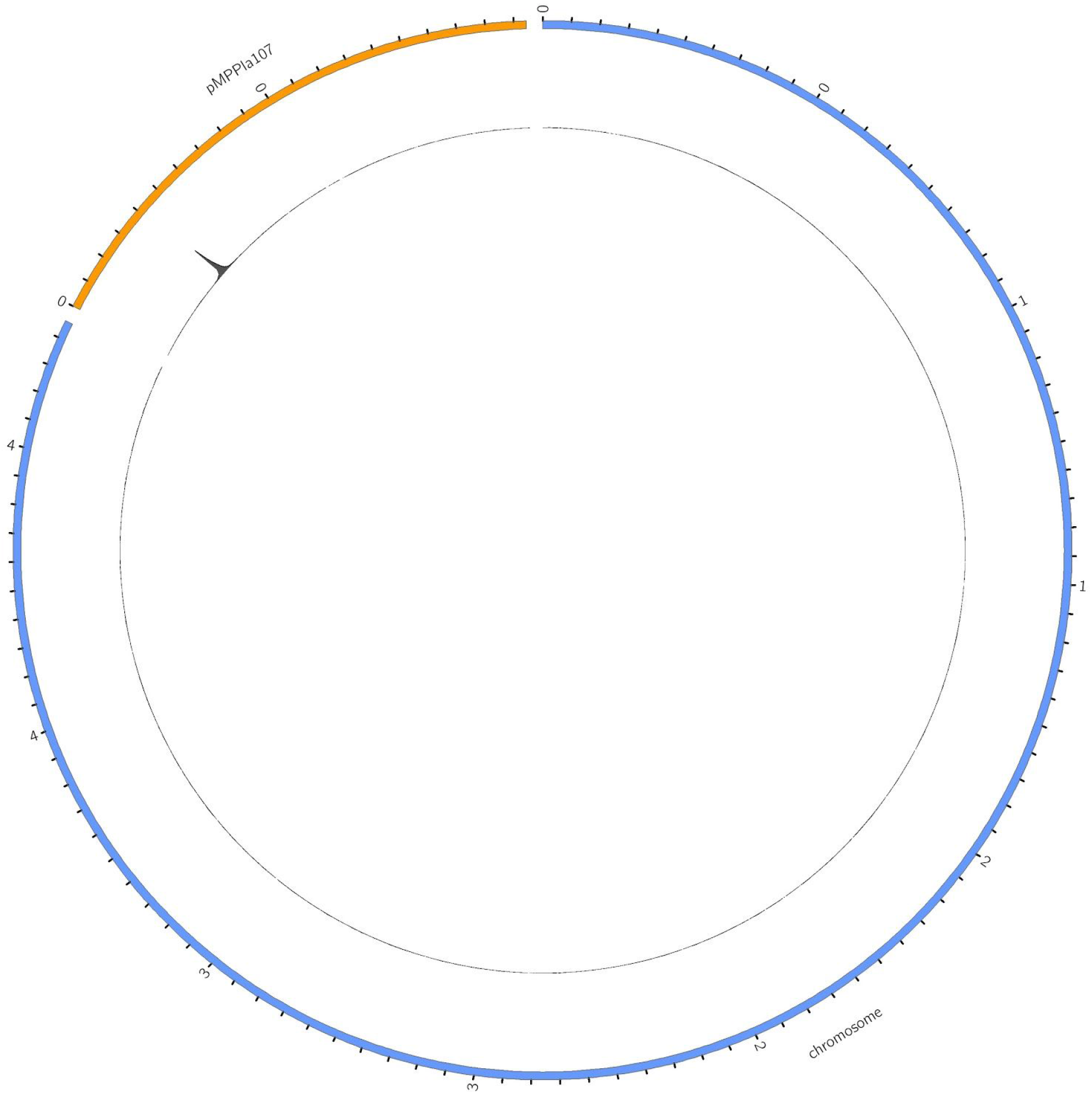
CRISPR-Cas Targeted Enrichment of a Transposon Insertion in *Pseudomonas stutzeri*. Insertion of a Mar2xT7 transposon into megaplasmid was pMPPla107 was created using a combination of selection for antibiotic resistance and screening for the ability of antibiotic resistance to be mobilized through conjugation to recipient strains. Ultimately, this tagged version of megplasmid pMPPla107 was conjugated into a nalidixic acid resitant isolate of *Pseudomonas stutzeri* 28a24 to create strain DBL494. This figure shows the results of CRISPR-Cas enrichment followed by sequencing using an Oxford Nanopore MinION of strain DBL494. Individual reads were mapped to either the chromosome of *Pseudomonas stutzeri* (blue) or megaplasmid pMPPla107 (orange), and the resulting coverages were plotted on the innermost ring. There is a spike in coverage surrounding a single transposon insertion point.

### Strategies to Overcome Bottlenecks in Transposon Insertion Identification through Whole Genome Sequencing

Since one of the well known strengths of the Oxford Nanopore MinION is the ability to sequence long reads with relative ease, our main driving question was not necessarily whether transposon insertions sites could be captured by single reads. Rather, our primary motivation for these experiments was to vet the ability of different sequencing kits and multiplexing options in order to find the optimal strategies to identify these insertions in a cost and time efficient way. A clear take away from our data is that the main cost bottlenecks in using Nanopore reads to identify transposon insertion sites are current limitations in multiplexing the reads and the sequencing depth available on single flowcells. In every case we were able to sequence genomes with adequate coverage to identify at least one transposon insertion site, however, a far majority of the reads using the RAPD sequencing kit were not able to be assigned to particular genomes and were thus tossed into a pile of reads with unknown origins. If reads were to be better demultiplexed, it would certainly cut down on the amount of sequencing depth required to have a good chance of identifying insertion sites. Moreover, use of this kit is currently further limited because only 12 barcodes are currently available. On the other hand, we note that we recovered a significantly higher percentage of reads from the ligation based kits and there are currently 24 barcodes available, and as such suggest use of these kits at present to maximize data generation.

Even in the absence of better software or multiplexing options, there are still experimental ways to maximize the recovery and assignment of transposon insertions to different mutants. Since the whole genome sequencing strategy is agnostic to the particular transposon used, so long as one knows the particular sequence to search for, multiple transposons or even different versions of each transposon could be used within single genomes and then sequenced together. In this case, the particular mutants would in effect be “barcoded” by the sequences of the transposon within them. Such a strategy would allow one to stack similar genomes containing different flavors of transposon into a single barcode and would thus allow for sequencing even more genomes than current multiplex options allow. Likewise, if the same transposon is used for genomes from strains that differ enough in genome sequence, these genomes could also be stacked into a single barcode or sequencing library and would be able to be identified in the absence of particular barcodes, thereby allowing additional multiplexing.

### Efficient Enrichment of Transposon Insertion Sites through CRISPR-Cas

One of the downsides, both in terms of throughput and in terms of cost, of using whole genome sequences to identify transposon insertions is the amount of uninformative background genomic “noise” that gets sequenced as part of either the transposase or the ligation based protocols described above. Recently, CRISPR-Cas strategies have been introduced that enable one to selectively sequence specific regions of the genome so long as crRNA can be developed to target those regions. In this case, transposon insertions are the perfect scenario for using targeted sequencing methods because the transposon sequence doesn’t change regardless of where it inserts. Therefore, one can design sets of crRNA that specifically target the transposon sequence and simply sequence out to identify the insertion points. As we show in Figure 1, this strategy is highly efficient for identifying transposon insertion points, as a far majority of reads that we sequenced as part of this method matched the region of the genome bracketed by the Mar2xT7 transposon. This technique significantly cuts down on the background level of genomic noise that needs to be sorted through to identify transposon insertions, and thus it is likely that many more libraries could be run on a single flowcell using the CRISPR-Cas methods compared to whole genome sequencing for insertion identification. Furthermore, in this case, we have focused on a proof of principle experiment whereby one insertion site is identified, but there is no reason that this protocol can’t be altered to use barcodes and to multiplex different mutants per flowcell run. Moreover, this approach should scale up quite well to enable techniques such as TnSeq using an Oxford Nanopore MinION or to identify natural polymorphisms in transposon insertion sites. Extending this strategy to further lower the cost, it is possible to multiplex different crRNA targets within a single reaction and thus multiple transposons could be easily targeted during the creation of a single library. It is quite likely that this protocol can be further extended to identify other conserved insertions within genomes of interest, such as t-DNA insertions in plants.

### Overall Recommendations and Cost Considerations

The RAPD kit is by far the easiest and most straightforward methods of preparing DNA for sequencing using a MinION. However, in our hands for these experiments, it was also clear that a significant percentage of reads were lost from the RAPD kit libraries due to the inability to accurately demultiplex. We do not know if our results presented here on demultiplexing and the RAPD kit are generalizable, although we have seen similar results across other libraries, but this loss of information could significantly impact the ability to call transposon integration sites with a reasonable amount of sequencing depth. To this point, reads arising from the LSK109 kit were much more accurately demultiplexed, which resulted in many more informative reads per library than the RAPD kit, even though roughly the same amount of DNA was sequenced using each method.

For identifying particular transposon insertion sites, it is clear that the CRISPR-Cas based strategy performs much better than either the ligation or transposon based kits in terms of sequencing depth of the target region per sample recovered. In our experience here, you will have enough information within minutes of starting to sequence the CRISPR-Cas enriched library to be able to identify particular insertion points for a particular integration event. Since sequencing reactions can be stopped and the flowcell cleaned/reloaded with different DNA, and since one of the main determinants of flowcell life is the amount of DNA sequenced through it, the ability to quickly identify insertion points using this method provides numerous technical advantages over whole-genome based strategies. Simply put, many more transposon insertions can be screened on the same flowcell using CRISPR-Cas enrichment because identification of insertions within each particular strain can occur without wasting sequencing depth on non-target genomic regions. Likewise, background sequencing noise increases with genome size, so targeted enrichment would likely be much more efficient than whole genome sequencing when trying to identify transposon or t-DNA insertions in organisms like plants or animals. However, the CRISPR-Cas based strategy is also the most expensive of the three methods explored here, at least on a reagent basis, because at baseline it requires the same materials as ligation based strategies in addition to oligos and CRISPR-Cas enzymes. This expenditure is somewhat overcome by volume since ordered oligos can be used for many independent reactions and can be pooled in single reactions against different targets. We also highlight that our current strategy used 4 different oligos to identify the Mar2xT7 transposon insertion, but that lowering this number to 3 or 2 oligos could lower the overall costs. Lastly, we note that while this method appears to work quite well for the identification of single transposon insertions, using CRISPR-Cas enrichment to identify insertions across a much broader library (as per TnSeq), will require the development of algorithms to identify true insertion points from random genomic noise generated by this method.

All of the experiments within this report involve a FLO-MIN106 flowcell, which currently costs upwards of 1000$/flowcell but which also generally enables sequencing of 10+Gb/flowcell. Although this cost and the chance at contamination across sequencing rungs is significantly cut through the use of protocols designed to wash and reuse a single flowcell as well as employment of nuclease flushes, the ability to sequence and identify transposon insertions using the ~90$ Flongle will certainly the process more cost efficient per strain. Although the Flongle currently enables roughly ¼ of the sequencing depth of the larger MinION flowcells, our experiments demonstrate that this level of depth should be more than adequate for identifying transposon insertions and especially through the use of targeted enrichment.

### Conclusions

Here we have demonstrated that long reads arising from the Oxford Nanopore MinION can be used to accurately identify transposon insertion sites within bacterial genomes. We have sequenced two different transposons across a variety of Gram negative bacterial strains, and in each case were able to identify insertion points as long as there was at least 2x coverage of the genome. From these results, it is our opinion that the LSK109 ligation based kits yield the best value when using MinION sequencing to identify transposon insertion points because the extra time that it takes compared to the RAPD kit is more than made up for in the amount of useable data that is generated. Furthermore, if cost is no issue, CRISPR-Cas based targeted sequencing provides a further boost in the signal to noise ratio for identifying transposon insertions in bacterial genomes and will likely be increasingly useful for reverse genetics experiments in organisms as the overall genome size increases.

## Acknowledgements

Many thanks to Nick Loman, Jennifer Gardy, Josh Quick, Jared Simpson, Matt Loose, John Tyson, Lauren Cowley and the whole PoreCamp Vancouver crew for providing a great environment to learn about Nanopore sequencing. Thank you also to Mark Martin for providing *Ensifer adhaerens*, and Victor de Lorenzo for providing a Tn7: *xylR* promoter -> *lacZ* version of the *Pseudomonas putida* strain KT2442.

